# Microcephaly-like phenotype triggered by novel reassortant and prototypic Oropouche Virus strains in brain organoids

**DOI:** 10.1101/2025.08.13.669874

**Authors:** Gabrielle Brum Lopes da Silva, Vivian Grizente Rocha, Fábio Luís Lima Monteiro, Beatriz Luzia de Mello Lima Guimarães, Livia Goto-Silva, João Marcos de Azevedo Delou, Pedro Junior Pinheiro Mourão, Ismael Carlos da Silva Gomes, Enzo Oliveira Barone, Matheus Villanueva Andrade, Rafael Ferreira Lima, Izabela Mamede, Clarisse Rezende Reis, Fernanda Martins Marim, Victor Emmanuel Viana Geddes, Renato Santana Aguiar, Amilcar Tanuri, Carolina Moreira Voloch, Stevens Kastrup Rehen

**Author notes:** These authors contributed equally to this work (Gabrielle Brum Lopes da Silva, Vivian Grizente Rocha, Fábio Luís Lima Monteiro).

## Abstract

Oropouche virus (OROV) is an emerging arbovirus currently spreading across South America, with increasing reports of neurological manifestations, severe systemic disease, and congenital abnormalities. Although traditionally associated with mild febrile illness, the recent geographic expansion and surge in OROV outbreaks have prompted attention to its neurotropic potential. Here, we investigated the impact of OROV infection on human neural development using neural stem cells (NSCs) and brain organoids derived from induced pluripotent stem cells. Recent OROV isolates exhibiting genomic reassortment and associated with increased neurological manifestations were compared with a prototypical strain for the ability to infect NSCs, early-stage organoids, and more mature cortical-like tissues. OROV infected NSCs efficiently, leading to widespread cell death, depletion of proliferative progenitors, and disruption of neuroepithelial organization. Transcriptomic profiling of infected NSCs revealed a robust reduction of antiviral response genes and an enrichment of pathways related to viral replication, apoptosis, and the inhibition of stem cell maintenance and neuronal differentiation. These molecular signatures aligned with the phenotypic collapse of progenitor pools and cortical structure observed in organoids. OROV antigens were detected in both astrocytes and neurons, with associated structural degeneration. Although a substantial overlap in differentially expressed genes was observed between the two viral strains, some strain-specific transcriptional responses were detected. However, these modest differences did not translate into distinct cytopathogenic effects between the two viral strains. These phenotypes, including the reduced growth of infected organoids, resemble those previously described with Zika virus in the same cellular models, supporting the hypothesis that OROV may impair brain development. Together, these results reveal a previously unrecognized neuroteratogenic potential of OROV strains and provide mechanistic insight into the potential of OROV to induce microcephaly-like phenotypes, highlighting its relevance as a significant threat to maternal-fetal health.

## Introduction

Oropouche virus (OROV) is an emerging arbovirus of the *Orthobunyavirus* genus (*Peribunyaviridae* family) that has historically caused self-limiting febrile illness in South America^1^. Recently, increasing reports of encephalitis, multiorgan failure, and notably, vertical transmission, have prompted urgent investigation into OROV neuropathogenesis.

In 2024, a new wave of outbreaks marked a shift in the epidemiological profile of OROV. Autochthonous transmission reported in Central America and southeastern Brazil indicated ongoing geographic expansion, likely supported by local ecological conditions^2,3^. Genomic analyses have identified reassortment events with OROV strains circulating in the South American Amazon region associated with these outbreaks^4^, with preliminary *in vitro* analysis suggesting altered replication and virulence^5^. In more than one fatal case, OROV antigens were detected in multiple organs, including the brain, and were associated with severe tissue inflammation and immune dysregulation^6^. Noteworthy, a case series documented OROV RNA and antigen in fetal tissues of newborns with microcephaly, reinforcing the possibility of vertical transmission^7^.

Understanding how OROV interacts with neural stem cells (NSCs) and differentiated cells is crucial, particularly in light of its similarities with Zika virus (ZIKV), which causes a major congenital syndrome characterized by cortical dysgenesis and microcephaly^8,9^. While animal models and adult human brain slice cultures have shown that OROV infects neurons and microglia and induces an inflammatory response, little is known about the virus’s effects during the early stages of human brain development^10–12^. Human induced pluripotent stem cell (hiPSC)-derived brain organoids recapitulate key aspects of fetal brain development, including neural progenitor proliferation, differentiation, and cortical organization, and have become valuable platforms for modeling congenital viral infections^13–15^. These three-dimensional systems enable the simultaneous assessment of viral tropism, cytopathic effects, and disruptions in neurogenesis^16^, offering a physiologically relevant context to investigate neurodevelopmental consequences of viral exposure.

Despite increasing reports of OROV circulation and its potential to reach new geographic regions, its impact on the developing human brain remains poorly characterized. So far, no study has investigated OROV infection in human NSCs or brain organoids. To address this gap, we employed hiPSC-derived neural stem cells (NSCs) and brain organoids to assess the infectivity and neuropathogenic potential of two genetically distinct OROV strains, including a recent clinical isolate exhibiting genomic reassortment.

To our knowledge, this is the first study to investigate OROV infection in human brain organoids, providing direct evidence of its neurodevelopmental impact using a physiologically relevant 3D *in vitro* model. We show that OROV infects NSCs, astrocytes, and neurons, leading to increased cell death, impaired proliferation, and disruption of cortical-like tissue organization. Transcriptomic analyses revealed that OROV alters neurodevelopmental programs in neural progenitors, consistent with the structural abnormalities observed in infected organoids. Together, these findings provide mechanistic insight into OROV-induced neurodevelopmental pathology and highlight its potential to contribute to congenital disease, particularly in the context of expanding viral transmission.

## Methods

### Virus strains and propagation

Two Oropouche virus (OROV) strains were used: BeAn19991 and LVM-2024. BeAn19991 is a reference strain obtained from the Evandro Chagas Institute, Brazil (GenBank accession numbers: L: KP052850; M: KP052851; S: KP052852). LVM-2024 is a clinical isolate collected in 2024 from the Cacaria district, Rio de Janeiro state, Brazil (GenBank accession numbers: L: PQ537324; M: PQ537328; S: PQ537332)^5^. These isolate displays segmental reassortment in the L and S segments, showing phylogenetic similarity to Iquitos virus (IQTV), an orthobunyavirus associated with outbreaks in Peru. Viral stocks were propagated in *Aedes albopictus* C6/36 cells (ATCC CRL-1660) maintained in Leibovitz’s L-15 (Gibco) medium supplemented with 2% fetal bovine serum (Gibco), non-essential amino acids (Merck), and 0.3% tryptose phosphate broth (Gibco), grown at a density of 7×10^4^ cells per cm^2^. Infections were performed at a multiplicity of infection (MOI) of 0.005. After 72h of infection at 28 °C, supernatants were collected, centrifuged at 300 × g for 10 min, filtered through a 0.22 µm membrane, aliquoted, and stored at -80 °C. Conditioned medium from uninfected C6/36 cells was used as mock control.

### Human iPSC culture and induction of Neural Stem Cells

Human induced pluripotent stem cells (hiPSCs) were maintained on Geltrex-coated plates in StemFlex medium (Thermo Fisher Scientific) under standard conditions (37°C, 5% CO_2_). hiPSCs were differentiated into NSCs according to the PSC Neural Induction Medium protocol (Gibco). Briefly, iPSCs were plated as cell clumps at a density of 2.5-3×10^5^ cells per well on Geltrex matrix-coated 6-well plates. On day 1, the culture medium was substituted with PSC Neural Induction Medium (Neurobasal plus 1X Neural Induction Supplement - NIS). Medium changes were performed every other day. By day 7, P0 NSCs were harvested and subsequently expanded in Neural Expansion Medium (1:1, Neurobasal:Advanced DMEM/F12, 1X NIS).

### Neural Stem Cell infection

Neural Stem Cells were cultured on Geltrex-coated plates for infection assays with OROV strains BeAn19991 or RJ (LVM-2024) at a multiplicity of infection (MOI) of 0.1. Cells were incubated for 2 hours with the viral inoculum at 37°C. Then, the medium was removed and replaced with fresh Neural Expansion Medium. For immunofluorescence analysis, cells were seeded in 96-well plates at a density of 1×10^4^ cells per well. Immunofluorescence analysis was performed in cells fixed at 24 hours post-infection (hpi). Cells were rinsed three times with phosphate-buffered saline (PBS), fixed in 4% paraformaldehyde (PFA) for 15 minutes at room temperature, and stored at 4°C until further processing.

For transcriptomic analysis, cells were plated in 24-well plates at 1×10^5^ cells per well. After 24 h of infection, conditioned medium was discarded, cells were washed twice with PBS and RNA was extracted.

### Synthetic RNA standard curve

To enable accurate RNA quantification in supernatants from infected NSCs, synthetic viral RNA produced via in vitro transcription was used to construct a standard curve. Initially, viral RNA was purified from OROV LVM-2024 stock using the RNeasy Mini Kit (QIAGEN), according to the manufacturer’s protocol. This RNA was reverse transcribed with Superscript IV (Thermo Fisher Scientific) using specific primers targeting the S segment (BstBI F: 5’ TGTCTTCGAAAGTAGTGTGCTCCACAATTCA 3’; XhoI R: 5’ GTCTTCCTCGAGAGTAGTGTGCTCCACTATATGTCA 3’). Following cDNA synthesis, the S segment was amplified using the Platinum II Taq Hot-Start DNA Polymerase system (Thermo Fisher Scientific). The amplicon was cloned into a TVT7R(0,0)^17^ vector (AddGene) via restriction digestion and ligated using T4 DNA Ligase. The resulting plasmid, designated TVT7OROV-S, was used as the template for an in vitro transcription reaction with the MEGAscript™ T7 Transcription Kit (Thermo Fisher Scientific), following the manufacturer’s instructions. Transcribed RNA was purified by lithium chloride precipitation and quantified using the Qubit™ RNA BR Assay Kit (Thermo Fisher Scientific). The synthetic RNA was then diluted to 2×10^8^ copies/µL, yielding 10^9^ copies per RT-qPCR reaction, followed by a serial 10-fold dilution (from 10^9^ to 10^2^ copies) in RNase-free water. All primers used were obtained from Exxtend.

### Viral load quantification

To assess the viral load in the supernatant of OROV-infected NSCs, RT-qPCR was performed in parallel with a synthetic RNA standard curve. Briefly, supernatants were collected 24 hours post-infection, and nucleic acids were extracted using the RNeasy Mini Kit (QIAGEN), following the manufacturer’s protocol. Purified RNAs from both the supernatants and synthetic RNA were then amplified using the GoTaq® Probe 1-Step RT-qPCR System, targeting the OROV S segment (OROV_FNF: 5’ TCCGGAGGCAGCATATGTG 3’; OROV_FNR: 5’ ACAACACCAGCATTGAGCACTT 3’; OROV_FNP: 5’(FAM) CATTTGAAGCTAGATACGG 3’)^18^. Following amplification, cycle threshold (Ct) values were plotted against the synthetic RNA standard curve to determine the corresponding RNA copy number in each sample using a simple linear regression method with GraphPad Prism 9.

### RNA extraction, quantification, library preparation, RNA-sequencing and analysis

Cell monolayers were incubated with Buffer RLT from the RNeasy Mini kit (Qiagen), and total RNA was extracted following the manufacturer’s procedures. Purified RNA was treated with TURBO DNA-free™ DNase to remove genomic contaminating DNA (ThermoFisher Scientific). Total RNA was quantified using Qubit RNA HS Assay Kit on a Qubit 4 fluorometer (ThermoFisher Scientific), and RNA integrity was assessed by capillary electrophoresis on a 4150 TapeStation System using a High Sensitivity RNA ScreenTape (Agilent Technologies). Only samples with a RNA Integrity Number (RIN) of ≥ 8.7 were used. Sequencing libraries were prepared from 100 ng of total DNase-treated RNA from 5 replicates from 2 independent experiments for each group (non-infected cells, OROV BeAn19991 infected cells, or OROV RJ LVM-2024 infected cells) using Illumina Stranded Total RNA Prep, Ligation kit (Illumina) according to the manufacturer’s instructions. Ribosomal RNA was depleted by the Ribo-Zero Plus kit, followed by fragmentation and denaturation of RNA, synthesis of first and second cDNA strands, adenylation of 3’ ends, anchor ligation, cleanup of fragments, library amplification and cleanup. For the cleanup steps, Agencourt RNAClean XP and Agencourt AMPure XP magnetic beads (Beckman Coulter) were used for RNA and for the libraries, respectively. Samples were indexed using IDT for Illumina DNA/RNA UD Indexes Set A. The prepared libraries were quantified using QIAseq Library Quant Assay kit (Qiagen), and the average size (350 bp) was verified using High Sensitivity D1000 ScreenTape reagents (Agilent Technologies). Sequencing was performed as paired-end using a P2 Reagent Cartridge 200 Cycles on the NextSeq 1000 platform (Illumina), with over 15 million reads per sample. Raw FASTQ files were quality-checked using FastQC and aligned to the human transcriptome (GENCODE v48) with Salmon^19^, applying default parameters and the Gibbs sampling bootstrap method. Transcript-level quantifications were imported into R and summarized at the gene level. All downstream analyses and visualizations were performed entirely in R, with the full code available at: https://github.com/iza-mcac/2025-08-Oropouche-virus-strains-brain.

### Brain organoids generation and culture

Brain organoids were generated from hiPSCs cultured in StemFlex medium on Geltrex-coated plates using an adapted version of the unguided Lancaster protocol^13,20^. Embryoid bodies (EBs) were formed by plating 9000 cells/well in 96-well plates in StemFlex medium containing 50 µM ROCK inhibitor (Merck Millipore) and centrifuging to improve initial aggregation. After day 1, EBs were cultured with human Embryonic Stem Cell medium (hESC medium): DMEM/F12, 20% Knockout serum replacement (KSR), 3% certified fetal bovine serum (FBS), 1% Penicillin-Streptomycin (P/S), 1% GlutaMAX (Life Technologies), 1% MEM-NEAAs and 100 µM 2-Mercaptoethanol (Thermo Fisher Scientific), 50 µM ROCKi, and 4 ng/mL bFGF (Thermo Fisher Scientific). On day 6, EBs were transferred to 24-well ultralow-attachment plates with neuroinduction medium: 1% N2 supplement (Gibco), 1% GlutaMAX, 1% MEM-NEAAs, 1% P/S (Thermo Fisher Scientific), and 1 µg/ml heparin (STEMCELL Technologies) in DMEM/F12. At day 10, organoids were coated with Matrigel (Corning) in a 60-mm non-adherent tissue culture plate. Following Matrigel coating, organoids were returned to 24-well ultra-low-attachment plates with neurodifferentiation medium without vitamin A: 50% neurobasal medium, 0.5% N2, 1% B27 supplement without vitamin A, 50 uM 2-Mercaptoethanol, 0.5% MEM-NEAA, 1% GlutaMAX, 1% P/S, and DMEM/F12 (Thermo Fisher Scientific) with media change after 48h in culture. On day 15, brain organoids were transferred to 6-well plates on an orbital shaker at 90 rpm and transferred to neurodifferentiation medium with vitamin A (Thermo Fisher Scientific). Medium was replenished twice a week until day 30 or 60.

### Brain organoid infection

Human brain organoids cultivated for 30 and 60 days in culture were infected with 3 × 10^3^ plaque-forming units (PFU) of OROV (strains BeAn19991 or RJ LVM-2024) per organoid for 2 h at 37°C under continuous agitation (90 rpm). Infections were performed in 12-well low-attachment plates containing 800 µL of neurodifferentiation medium with vitamin A per well. After infection, organoids were washed three times with PBS and cultured in virus-free differentiation medium under continuous agitation. Conditioned media from organoids at 30 days of differentiation were collected at 12 h and 1 to 5 days post-infection (dpi) for LDH assays (Promega), 10 µL, and viral titration (50 µL). At 5 dpi, organoids were fixed in 4% paraformaldehyde (PFA) for 1 h at room temperature, washed with PBS, cryoprotected in 30% sucrose and stored for cryosectioning.

Organoids at 60 days in culture were maintained until 11 dpi with medium changes (neurodifferentiation medium with vitamin A) on days 3 and 7 post-infection. Endpoint analysis was performed at 11 dpi, when organoids were fixed in 4% paraformaldehyde (PFA) for 1 h at room temperature, washed with PBS, cryoprotected in 30% sucrose and stored for cryosectioning.

### Viral titration

Viral titers were measured by plaque assay in Vero Cells as described previously^5^. Briefly, Vero CCL-81 cells (BCRJ-0245) seeded in 24-well plates at 90% confluency were inoculated with 10-fold serial dilutions of OROV stocks or organoids supernatant for 1 hour in the incubator (37 °C, 5% CO2). After viral adsorption, the inoculum was removed, and cells were cultivated for 3 days in a semisolid medium composed of Alpha-MEM supplemented with 1% fetal bovine serum, 1% penicillin-streptomycin, and 1.5% carboxymethyl cellulose (Merck). After this period, cells were fixed with 10% formaldehyde solution for 1 hour, washed with distilled water and stained with 0.25% crystal violet and 20% ethanol solution for plaque visualization. Viral titers were expressed as plaque formation units (PFU) per milliliter. For all inoculations the MOI was calculated based on viral infectious units (PFU) per number of cells.

### Cytotoxicity assays

OROV-induced cytotoxicity was assessed by measuring lactate dehydrogenase (LDH) release into the culture medium as a marker of cell lysis. At each point, 10 µL of conditioned medium was collected and mixed with 90 µL of LDH storage buffer (200 mM Tris-HCl, pH 7.3; 10% glycerol; 1% BSA). Samples were stored at −80°C and thawed simultaneously for batch processing. LDH levels were quantified using the LDH-Glo™ Cytotoxicity Assay (Promega), according to the manufacturer’s protocol. For each condition, LDH values were normalized to the 12-hour post-infection time point to calculate fold changes.

### Immunofluorescence and image acquisition

NSC cultured in 96-well plates were fixed with 4% formaldehyde for 15 minutes at room temperature. After fixation, samples were washed for 5 min with PBS, permeabilized with 0.3% Triton X-100 in PBS for 15 minutes and washed again with PBS for 5 min. Blocking was performed for 1 hour at room temperature using 3% BSA and 5% NGS in PBS. Primary antibodies were diluted in blocking solution and incubated for 1 hour at 37°C. The following primary antibodies were used: anti-OROV (mouse polyclonal anti-OROV, ATCC, antibody, in-house, 1:1000) and anti-Nestin (rabbit, 1:500). Following primary antibody incubation, cells were washed three times for 5 min each with PBS, fluorescently labeled secondary antibodies (Alexa Fluor 488 donkey anti-mouse and Alexa Fluor 568 donkey anti-rabbit; Invitrogen, 1:400) were incubated for 2 hours at 37°C, protected from light. Nuclei were stained for 10 min with DAPI (300 nM solution in PBS), followed by two PBS washes and a final rinse with MilliQ water. A solution of 50% glycerol in PBS was added to the plates. Fluorescence images were acquired using an Agilent Cytation 1 Cell Imaging, and quantification was performed using Gen5’s integrated analysis tools.

Brain organoids at 30 and 60 days in culture were fixed in 4% paraformaldehyde for 1 hour at room temperature, rinsed with PBS, cryoprotected overnight in 30% sucrose at 4 °C, embedded in Tissue Freezing Medium (Leica). Cryosections of 10 μm were obtained using a Leica CM1860 cryostat. Antigen retrieval was performed for all conditions by heating sections in citrate buffer: 10 mM Sodium Citrate, 0.05% (v/v) Tween 20, pH 6.0. Organoid sections were permeabilized in 0.3% Triton X-100 in PBS for 15 minutes, washed in PBS for 5 minutes, and blocked for 1 hour in 3% BSA prior to overnight incubation at 4 °C with primary antibodies diluted in blocking buffer (3% BSA and 5% normal goat serum in PBS). The following primary antibodies were used: anti-cleaved caspase-3 (rabbit, Millipore, 1:100), anti-OROV (mouse, ATCC, 1:200), anti-PAX6 (rabbit, Thermo, 1:100), anti-Ki67 (rabbit, Neuromics, 1:100), anti-GFAP (rabbit, Abcam, 1:250), and anti-MAP2 (rabbit, Invitrogen, 1:100). After washing, sections were incubated with secondary antibodies (goat anti-rabbit or anti-mouse Alexa Fluor-conjugated, Invitrogen, 1:400), and nuclei were counterstained with 300 nM DAPI for 10 minutes. Slides were mounted with Aqua-Poly/Mount and stored at 4 °C after drying. Immunofluorescence images were acquired using either a Leica TCS SP8 confocal microscope or a Zeiss Cell Discoverer, with a 20× objective lens. Image analysis was performed using Fiji and ZEN Blue software.

### Statistical analysis

Replicates and number of organoids analyzed are described in the specific results sections and figures subtitles. Data are presented as the mean ± standard error of the mean (SEM). Statistical analyses were performed using GraphPad Prism 9. Comparisons between multiple groups were performed using one-way ANOVA followed by Tukey’s multiple comparison test. Differences were considered statistically significant when p < 0.05.

## Results

### OROV infects human NSCs and modulates neurodevelopmental markers

To assess the susceptibility of neural progenitor cells to OROV, we exposed human iPSC-derived NSCs to two viral strains: the prototypical BeAn19991 strain and a recently isolated clinical strain from Rio de Janeiro (RJ, LVM-2024) presenting genomic reassortment in the S and L segments associated with the recent virus expansion outbreaks in Latin America^5^. At 24 hpi, immunofluorescence staining revealed robust intracellular accumulation of OROV antigens (green) in Nestin-positive NSCs (red), confirming infection of both strains (Figure 1A). Notably, quantitative analysis at 24 hpi showed a higher percentage of infected NSCs in the BeAn19991 condition (86.6% ± 9.8%) compared to the RJ LVM-2024 strain (48.2% ± 8.8%), suggesting differences between strains (Figure 1B). In addition, both OROV lineages displayed detectable viral loads in the NSC supernatants (Figure 1C), reaching 3.39 × 10^4^ ± 2.07 × 10^4^ RNA copies per mL for BeAn19991 and 7.96 × 10^3^ ± 4.16 × 10^3^ for RJ LVM-2024 strain. These findings demonstrated that OROV efficiently infects human neural progenitor cells at early stages of differentiation and raises the possibility of interference with neurodevelopmental programs.

**Figure 1.**
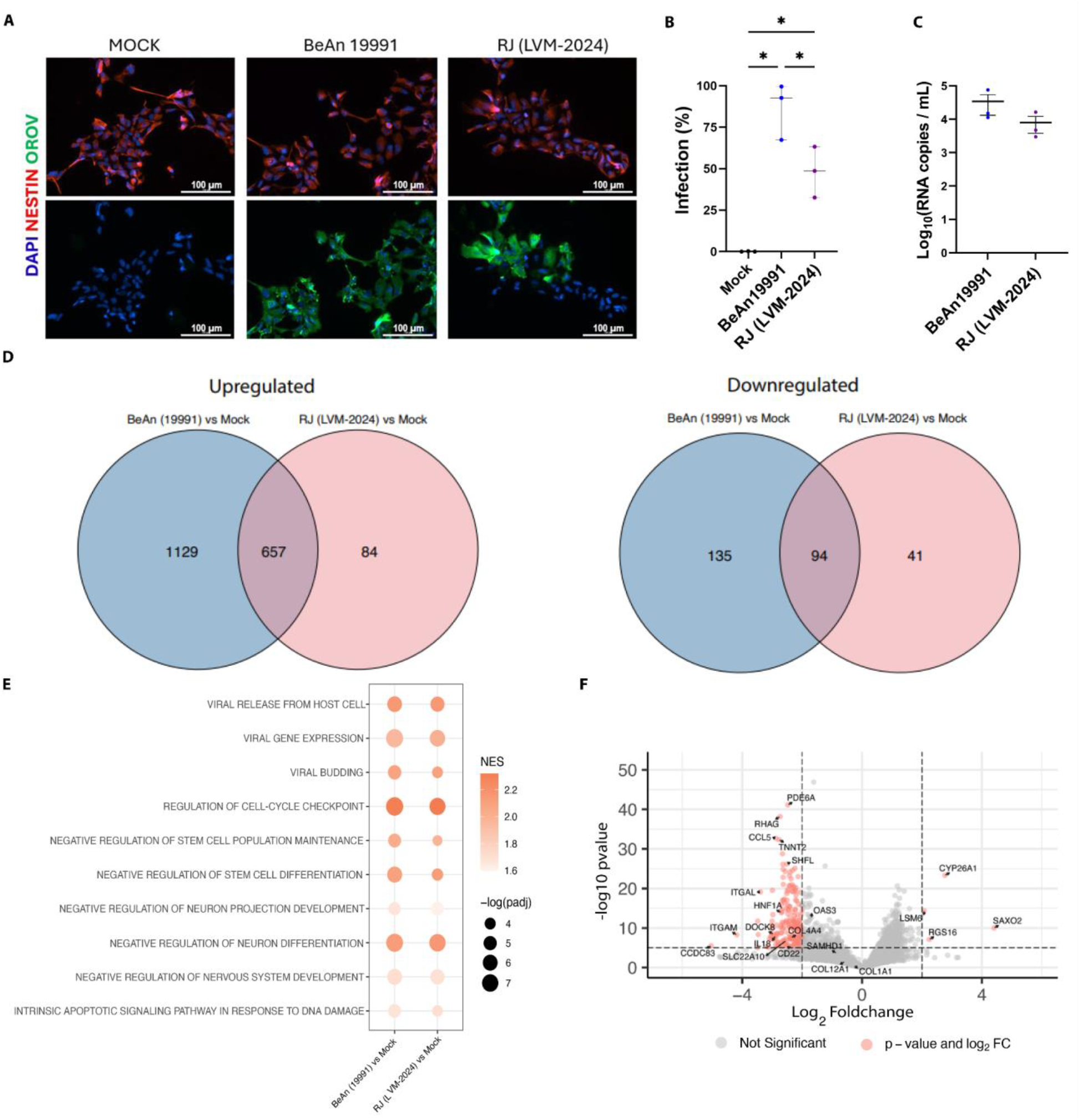
OROV infects human NSCs and modulates neurodevelopmental markers. (A-B) Immunofluorescence of hiPSC-derived NSCs at 24 hpi with OROV strains BeAn19991 or RJ (LVM-2024). Cells were stained with anti-OROV (green), anti-Nestin (red), and DAPI (blue). Both viral strains efficiently infected Nestin+ NSCs, with cytoplasmic accumulation of viral antigen. Scale bars = 100 µm. Data represent mean ± SEM from three independent experiments (^*^=*p*<0.05). (C) OROV RNA quantification in conditioned medium of NSCs infected at a MOI 0.1 for 24h. Data expressed as Log10 (RNA copies/mL) mean ± SEM of three independent experiments. (D) Venn diagrams showing differentially expressed genes (DEGs) in NSCs infected with BeAn19991 or RJ (LVM-2024) versus mock non-infected cells. (E) Gene Set Enrichment Analysis (GSEA) of shared DEGs revealed significant enrichment for pathways related to viral replication (e.g., “viral gene expression,” “viral budding”) and disruption of NSC function, including “negative regulation of stem cell population maintenance,” “negative regulation of neuron differentiation,” and “intrinsic apoptotic signaling.” NES: normalized enrichment score. Dot size reflects –log(padj). Data are representative of three independent transcriptomic experiments. (F) Volcano plot of RJ (LVM-2024) vs BeAn19991 comparison. Negative values represent BeAn19991 expression and positive values represent RJ (LVM-2024). Differentially expressed genes (absolute Log2FoldChange higher than 2 and p-value lower than 0.01) are colored in red.

To further characterize the molecular consequences of OROV infection in human NSCs, we performed transcriptomic analysis at 24 hpi. Differential gene expression analysis - applying thresholds of Log2FoldChange higher than 2 and p-value lower than 0.01 - revealed a large number of upregulated genes in both BeAn19991- and RJ (LVM-2024)- infected NSCs compared to mock controls, with 657 genes commonly induced across both strains (Figure 1D). Downregulated genes also showed substantial overlap (94 shared). Gene set enrichment analysis (GSEA) revealed regulation of pathways related to viral replication, such as viral gene RNA expression, viral release from the host cell and viral budding, as well as biological processes critical for early neurodevelopment. Both strains showed enrichment for gene sets associated with inhibition of stem cell maintenance and differentiation, and negative regulation of neuron projection development and neuronal differentiation (Figure 1E). Additionally, we observed enrichment of pathways related to the regulation of cell-cycle checkpoints and the intrinsic apoptotic signaling pathway in response to DNA damage, suggesting that OROV infection may trigger cell death mechanisms in NSCs (Figure 1E). These results provide molecular evidence that OROV impairs fundamental programs of neural progenitor function.

The comparison between BeAn19991- and RJ (LVM-2024)-infected NSCs (Figure 1G), revealed only 4 genes upregulated in RJ (LVM-2024): *CYP26A1, LSM6, RSG16* and *SAXO2*. These genes are associated with pathways related to host defense, cellular stress responses, and structural remodeling. Interestingly, interferon-stimulated genes (ISGs) such as *IL18, CCL5, APOBEC3A, ITGAL* and *ITGAM* were downregulated in RJ (LVM-2024) compared to BeAn19991. These genes appear upregulated in BeAn19991-versus Mock, but not in RJ (LVM-2024) versus Mock (Supplementary Figure 1, Supplementary Table S1). This suggests that LVM-2024 strain (RJ) elicits a weaker interferon response in NSCs than the reference BeAn19991. Additionally, the collagen-encoding gene *COL4A4* was downregulated in RJ (LVM-2024)-infected NSCs, while *COL12A1* and *COL1A1* showed a trend toward lower expression. These genes are essential for the development of the brain and the blood-brain barrier and were also downregulated in brains of newborns with congenital Zika syndrome^21,22^.

### Oropouche virus strains permissively infect brain organoids, leading to impaired growth and tissue organization

Infection of neuroprogenitors and regulation of gene expression affecting the developmental program prompted us to investigate if OROV infection in brain tissue derived from iPSCs could reproduce some of the deleterious effects of *in vivo* infection. For that, 30-day organoids were infected with BeAn19991- and RJ (LVM-2024) and monitored for 5 dpi.

Viral progeny was quantified in supernatants at multiple time points after inoculation by plaque assay. Both strains replicated with detectable viral titers as early as 12 hpi (Figure 2A), with the BeAn19991 strain showing higher titers. After 24 hours, viral titers increased during infection with no significant differences between the strains. These data indicate that both OROV strains are capable of infecting and replicating efficiently in human brain organoids.

**Figure 2.**
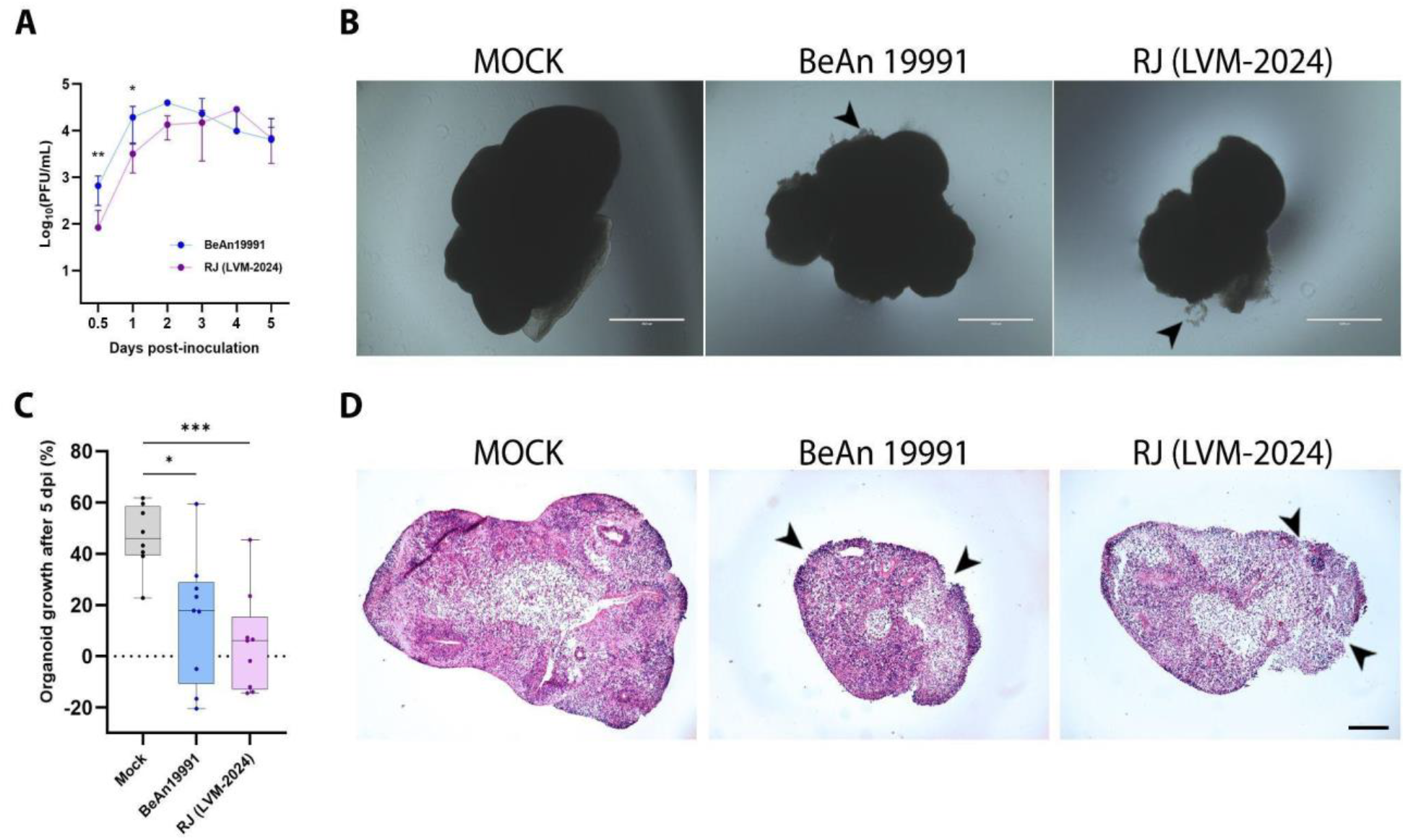
Oropouche virus strains permissively infect brain organoids, leading to impaired growth and tissue organization. (A) Viral production kinetics of OROV strains BeAn19991 and RJ (LVM-2024) in brain organoids. Viral titters were quantified in supernatants collected at multiple time points up to 5 dpi by plaque assay. Data are presented as mean ± SEM from two independent experiments, with at least 3 organoids in each condition. (B) Representative brightfield images of 30-day brain organoids at 5 dpi, after fixation. Scale bars = 1000 µm. (C) Quantification of brain organoid growth at 5 dpi, comparing organoids cultivated for 35 days to their size at day 30 of differentiation. OROV strains significantly reduced organoid growth compared to mock. Data represents SEM. (D) Hematoxylin and eosin (H&E) staining of 30-day brain organoids sections at 5 dpi reveals preserved structural organization in mock samples, characterized by intact radial arrangement and cellular density. In contrast, infection with BeAn19991 and RJ (LVM-2024) leads to marked tissue disorganization, including peripheral structural breakdown (arrowheads). Scale bars = 1000 µm.

Morphological inspection of infected organoids revealed reduced growth and irregular contours compared to mock controls (Figure 2B-C). Growth rate was reduced in both viral conditions when compared to mock treated cells, showing disruption of the development (Figure 2C). Histological sections revealed architectural disarray, with collapse of the neuroepithelial surface (Figure 2D). Cortical-like stratification, evident in controls, was severely disrupted in infected conditions. These observations indicate that OROV infection compromises both growth and structural organization of developing brain tissue.

### OROV infection induces widespread cell death in brain organoids

To determine whether cell death contributed to the observed collapse in organoid morphology, we performed cleaved caspase-3 immunostaining. At 5 dpi, both BeAn19991 and RJ (LVM-2024) strains triggered a marked increase in caspase-3 positive cells compared to mock (Figure 3A-C). Caspase-3 activation in infected organoids was not confined to superficial layers but extended throughout the neuroepithelium, including ventricular-like zones (Figure 3B). Supplementary Figure 2A-B shows the colocalization between OROV and caspase-3, indicating apoptosis in infected cells. Quantification (Supplementary Figure 2C) revealed no difference between strains, supporting comparable cytotoxic effects. These results suggest that OROV infection leads to widespread and strain-independent apoptosis in brain organoids.

**Figure 3.**
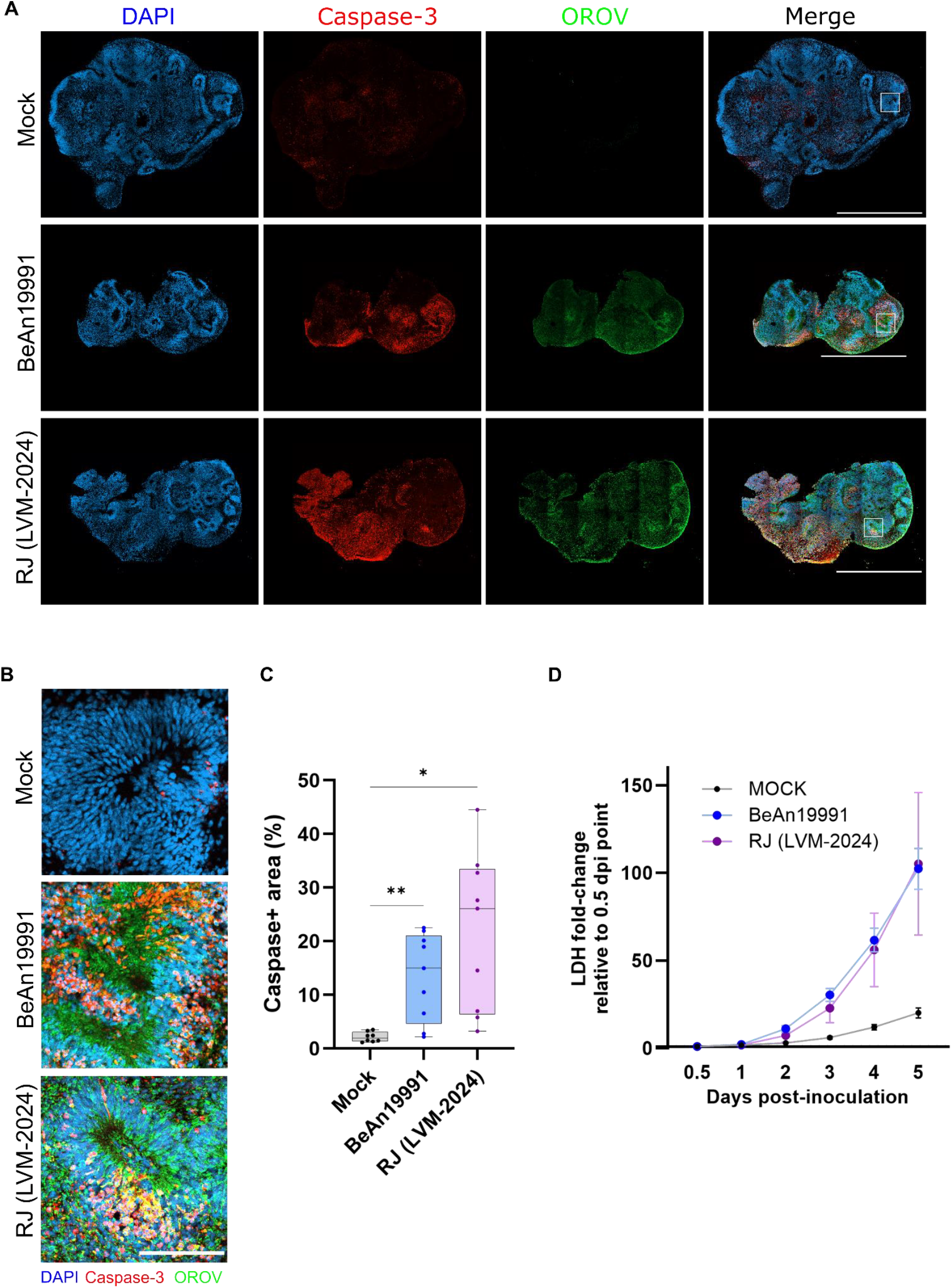
OROV infection induces widespread cell death in brain organoids. (A) Confocal images of 30-day brain organoid sections at 5 dpi with OROV strains BeAn19991 or RJ (LVM-2024), stained for DAPI (blue, nuclei), cleaved Caspase-3 (red, apoptotic cells), and OROV (green, viral protein). Scale bars = 1000 µm. (B) High-magnification confocal images of 30-day brain organoids at 5 dpi, from the insert area in (A). Scale bars = 100 µm. (C) Quantification of Caspase-3^+^ area relative to total organoid. Data represent mean ± SEM from two independent experiments, with at least 3 organoids in each condition (^*^=*p*<0.05; ^**^=*p*<0.01). (D) Cell death quantified by LDH release assay in 30-day organoid culture supernatants over 5 dpi. Data represent mean ± SEM from two independent experiments, with at least 3 organoids in each condition. Both strains significantly increased LDH release compared to mock, with no significant difference observed between the two viral strains.

Further evaluation for cell toxicity during the course of infection was obtained by monitoring the release of lactate dehydrogenase (LDH), a marker of cytotoxicity (Figure 3D). Along the 5 days of infection BeAn19991 and RJ (LVM-2024) induced a progressive increase in LDH release compared to the mock controls. No statistically significant differences were observed between strains at any time point, indicating comparable cell damage with similar kinetics. Altogether, these results support a comparable cytotoxicity profile for the two strains.

### Neural progenitor proliferation and identity are disrupted by OROV

Given the high infectivity in NSCs and the degenerative phenotype in organoids, we examined whether OROV impacts the neuroprogenitor pool in brain organoids. Immunostaining for Ki67 revealed a dramatic reduction in proliferating cells following infection (Figure 4A-B). Quantitative analysis revealed an almost complete loss of Ki67^+^ cells in ventricular-like zones, areas that during brain development contain organized neural progenitor cells. Similarly, the radial glial marker PAX6 was decreased and more disorganized in infected conditions (Figure 4C-D), with discontinuous expression along the apical surface. These results indicate that OROV infection depletes both the proliferative and structural neuroepithelial scaffold, likely contributing to impaired neurogenesis and collapse of tissue architecture.

**Figure 4.**
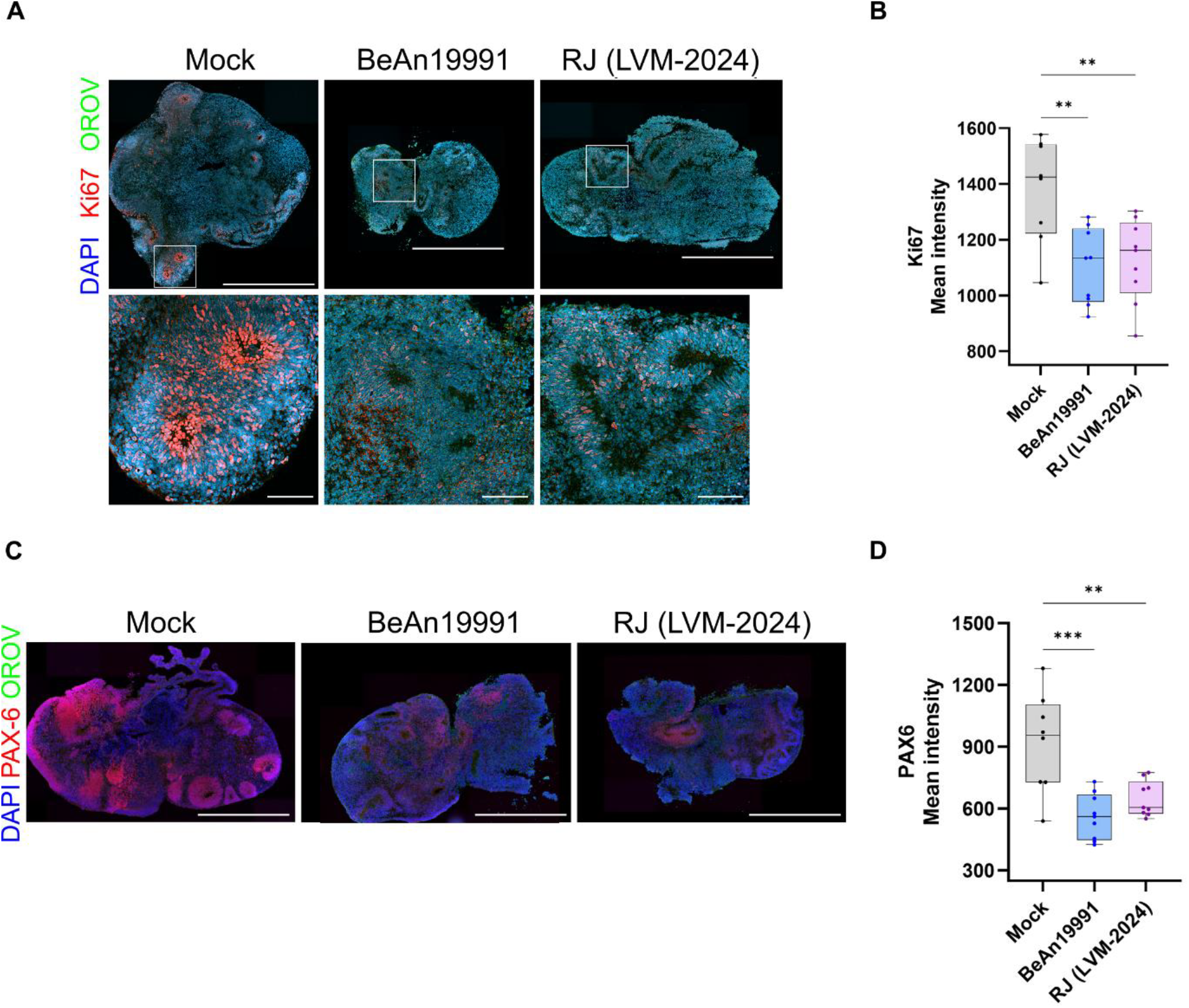
Neural progenitor proliferation and identity are disrupted by OROV. (A-B) 30-day brain organoid at 5 dpi with OROV strains BeAn19991 or RJ (LVM-2024), stained for DAPI (blue, nuclei), Ki67 (red, proliferating cells), and OROV (green, viral protein). Scale bars = 1000 µm. (B) Quantification of Ki67+ mean intensity shows a reduction in proliferative cells in infected samples compared to mock. No significant difference was observed between BeAn19991 and RJ (LVM-2024). Data represent mean ± SEM from two independent experiments, with at least 3 organoids in each condition. (C) 30-day brain organoid sections at 5 dpi with OROV strains BeAn19991 or RJ (LVM-2024), stained for DAPI (blue, nuclei), PAX6 (red, proliferating cells), and OROV (green, viral protein). Scale bars = 1000 µm. (D) Quantification of PAX6+ mean intensity shows a significant reduction in neural progenitor marker expression in infected samples compared to mock. No significant difference was observed between BeAn19991 and RJ (LVM-2024). Data represents a mean ± SEM from two independent experiments, with at least 3 organoids per condition.

### Oropouche virus infects astrocytes and neurons in brain organoids

To assess OROV tropism in neural tissues at later stages of development, we infected 60-day organoids containing differentiated astrocytes and neurons. Infection was carried out up to 11 days, when we confirmed citotoxicity by alterations in cytoarchitecture in infected tissue. While mock-infected organoids exhibited uniform cellular organization, those infected with the BeAn19991 and RJ (LVM-2024) strains displayed pronounced tissue disorganization, with extensive areas of cell death (Figure 5A).

**Figure 5.**
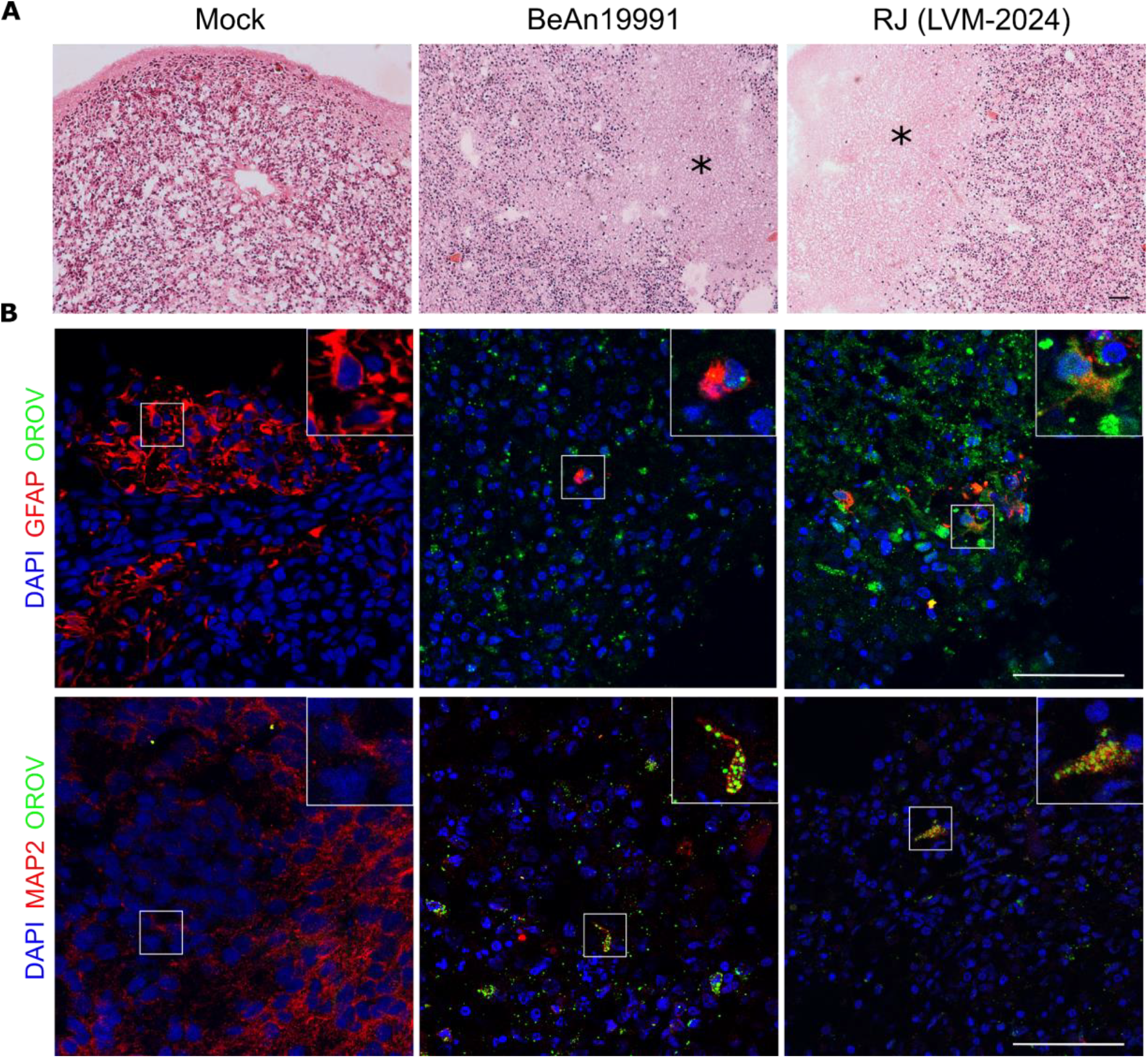
Oropouche virus infects astrocytes and neurons in brain organoids. (A) Brightfield images of hematoxylin and eosin-stained day 60 brain organoid sections at 11 dpi reveal substantial tissue damage in infected samples. While mock organoids show preserved cytoarchitecture, BeAn19991- and RJ-infected organoids display widespread tissue disorganization, including areas of cell death (indicated by asterisks) and disrupted structural integrity. Scale bar = 100 µm. (B) Brain organoids at 60-day were infected for 11 days and analyzed by immunofluorescence. Upper panels show sections stained for GFAP (astrocytes, red), OROV (green), and DAPI (nuclei, blue); lower panels show sections stained for MAP2 (neurons, red), OROV (green), and DAPI (blue). Images are representative of n = 3 organoids per condition. Scale bar = 100 µm.

Immunofluorescence revealed strong colocalization of OROV antigens with GFAP+ astrocytes and MAP2+ neurons (Figure 5B), indicating that OROV can infect not only neural progenitors but also differentiated neuronal and glial cells. These observations suggest that OROV infection extends beyond progenitors and contributes to global neural network degeneration.

## Discussion

Recent epidemiological data from the Brazilian Ministry of Health report five confirmed deaths resulting from OROV infection, with additional two fatalities currently under investigation. In 2024, approximately 13,000 cases were reported nationwide, and a steep rise is evident in 2025, with nearly 12,000 cases already documented by mid-year^23^.The aforementioned outbreaks were associated with a novel OROV reassortant lineage, raising concerns about changes in viral spread and pathogenesis associated with neurological complications, severe systemic disease, and suspected congenital infections^3,7,24,25^. In particular, a case series of newborns with microcephaly, arthrogryposis, and fetal demise associated with maternal OROV exposure resembled teratogenic effects observed in congenital Zika syndrome^12^, prompting the scientific community to investigate OROV’s impact on early development.

Pioneer work from our group showed microcephaly-like phenotype induced by ZIKV infection in human brain organoids^8^. Here, we show neuroepithelial collapse and reduced growth in brain organoids infected with two genetically distinct OROV strains, the prototype BeAn19991 and the recently circulating RJ (LVM-2024) isolate. The comparison of BeAn19991 and RJ (LVM-2024), revealed no major phenotypic differences in terms of cell death or tissue damage, although the prototypical strain displayed faster replication kinetics and higher titers at 12hpi. This contrasts with *in vitro* data suggesting increased virulence of reassortant strains circulating in Brazil^26^. While past reports raised concern over the potential emergence of more pathogenic variants, our findings do not support increased cytopathogenicity in neural models. Nonetheless, we should consider that more nuanced differences in viral behavior may only emerge when comparing RJ/LVM-2024 with other contemporary strains. In particular, comparative studies with the novel BR-2015-2024 reassortant lineage, identified during the 2023-2024 outbreaks in northern Brazil, may reveal subtle variations in tropism, replication dynamics, or immunoevasion^27^.

Martins et al. published a series of cases of malformations and miscarriages from 2015 to 2021 without definitive diagnoses which tested positive for anti-OROV IgM or viral RNA^12^, supporting the hypothesis of OROV vertical transmission. Specifically, a fatal case in a 47-day-old infant showing histopathological evidence of severe neuronal necrosis and widespread OROV antigen distribution, aligned closely with the organoid-level histology observed, thus reinforcing the translational relevance of our in vitro model. A recent study demonstrating that OROV is capable of infecting trophoblasts and human placental explants, contributed to the vertical transmission hypothesis^28^. Whether that investigation was limited to the BeAn19991 strain, a prototypical isolate from early Oropouche fever outbreaks, our study incorporates a representative strain from the current outbreak, when the first confirmed cases of congenital OROV infection were documented, and further demonstrates that reassortant OROV strains not only infect but also disrupt key neurodevelopmental processes. Collectively, these findings suggest that the teratogenic potential of OROV may be an intrinsic aspect of its pathogenesis, historically underdiagnosed due to its circulation in low-income regions such as the Amazon basin. Therefore, the recent surge in severe cases might reflect the broader geographic spread and higher incidence of OROV infection across Brazil, rather than intrinsic changes in viral virulence.

Transcriptomic profiling of infected NSCs reveals a direct impact of OROV on key neurodevelopmental programs. Both viral strains triggered upregulation of viral replication pathways and downregulation of gene sets involved in neural stem cell maintenance and neuronal differentiation. Notably, gene sets such as “negative regulation of neuron projection development” and “regulation of cell-cycle checkpoint” were significantly enriched, indicating that OROV impairs the molecular machinery responsible for proliferation and maturation of neural precursors. These molecular effects align with the collapse of neuroepithelial zones and loss of PAX6+ and Ki67+ cells observed in brain organoids, strengthening the mechanistic link between viral infection and disrupted corticogenesis. Moreover, infection by RJ (LVM-2024) strain downregulated the collagen-encoding gene *COL4A4* in infected NSCs and showed a trend toward lower expression of *COL12A1* and *COL1A1*, which are essential for brain development and the integrity of the blood–brain barrier^21,22^. Remarkably, previous studies of brains from neonates affected by congenital Zika syndrome reported a significant reduction in collagen fibers compared to control brains of the same gestational age^22^, reinforcing the potential teratogenicity of OROV strains.

OROV tropism for microglia and neurons and neuroinflammatory damage in adult human brain slice cultures was previously demonstrated by Almeida et al^29^. Our data expand these observations to the developing brain, demonstrating that NSCs are highly permissive to infection, with robust induction of cell death. In contrast to previously reported findings in adult tissue, showing negligible astrocyte infection^29^, we observed clear OROV tropism for both astrocytes and neurons in brain organoids. This suggests that the immature human brain may present a broader range of susceptible cell types, potentially contributing to the severity of congenital outcomes. Given the critical role of radial glia and astrocytes in scaffolding cortical development, the infection in these cells could have compounded effects on tissue organization and neuronal maturation. Noteworthy, while brain organoids used in the present work capture key features of early corticogenesis and viral pathogenesis, they lack a functional immune system and placental interface^30^. Future work incorporating microglia-containing assembloids or maternal–fetal interface models may better capture the interplay between innate immunity and viral neurotropism.

In conclusion, this study provides the first direct evidence that OROV infects and damages the human developing brain, impairing neurogenesis and tissue architecture, and thereby supporting its neurotropic and potentially teratogenic role. By leveraging hiPSC-derived brain organoids, we establish a mechanistic foundation to interpret congenital OROV cases and provide biological plausibility for its involvement in congenital disease. These findings highlight the need for heightened surveillance, inclusion of OROV in congenital infection diagnostic panels, and further studies exploring strain-specific virulence, innate immune responses, and long-term outcomes. Moreover, the in vitro platforms developed here set the stage for the investigation of antiviral agents, offering tools for a rapid response to future neurotropic arboviral outbreaks.

## Acknowledgements and Funding

We thank the contribution of the graduate students, Geovana Rodrigues Ferreira, Isabelle Cuba Teixeira Lopes, and Joana Cardoso do Prado Maciel, for technical assistance in performing immunostainings in brain organoids and NSCs. We thank Gabriel Ferraz da Silva for technical support with image acquisition at Zeiss Cell Discoverer and image quantification in Zen Blue software. We also thank Carolina Filgueiras Sanchez and Lendel Correia da Costa for their technical assistance with cell culture experiments. The authors acknowledge the use of AI-assisted tools (ChatGPT, OpenAI) for language refinement and editorial consistency during manuscript preparation. All scientific content and interpretation are the responsibility of the authors. All authors reviewed and approved the final manuscript before submission. This work was also supported by the following grants: Intramural grants provided by the D’Or Institute for Research and Education; Instituto Todos pela Saúde - ITpS (C1294; C2024-0779, A113; 27.909-9); Conselho Nacional de Desenvolvimento Científico e Tecnológico - CNPq (315592/2021-4, INCT- One CNPq 405786/2022-0); Fundação de Amparo à Pesquisa do Estado de Minas Gerais - FAPEMIG (APQ-02826-24); and Fundação Carlos Chagas de Amparo à Pesquisa do Estado do Rio de Janeiro - FAPERJ (E26/210.773-2021; E-26/201.356/2022; E-26/200.534/2025).

## Data Availability Statement

Transcriptomic raw data is available under project accession PRJNA1248042 in SRA ncbi.

## Author Contributions

S.K.R. conceived, designed, and coordinated the entire study. G.B.L.S. designed, planned, and performed all experiments, including the culture of human neural stem cells and brain organoids, as well as immunostaining procedures, interpreted the results, prepared figures, and wrote the manuscript.

V.G.R. designed, planned, and performed all experiments, including the culture of human neural stem cells and brain organoids, as well as immunostaining procedures, interpreted the results, prepared figures, and wrote the manuscript.

B.L.M.L.G. designed and planned experiments, including the culture of human neural stem cells and brain organoids, as well as immunostaining procedures, interpreted the results, prepared figures, and wrote the manuscript.

L.G.S. designed and planned experiments, discussed results, prepared figures, and performed immunostaining procedures, confocal image acquisition, quantification, data analysis, and reviewed the manuscript.

J.M.A. discussed results, prepared figures, performed image acquisition, quantification, data analysis and reviewed the manuscript.

I.C.S.G. conducted histological processing (embedding, sectioning, H&E staining) of all brain organoids, performed image acquisition, and prepared figures.

I.M., C.R.R., F.M.M., V.E.V.G., and R.S.A. conducted the RNA-seq analysis, discussed the results, prepared the figures, wrote and reviewed the manuscript.

F.L.L.M. designed, planned, and performed experiments, including viral inoculation and titration, cell viability assays, data curation and analysis; wrote and reviewed the manuscript.

P.J.P.M. and M.V.A. conducted experiments involving viral inoculation and titration, cell viability assessment, and sample collection.

E.O.B. and R.F.L. performed experiments including sample collection, nucleic acid extraction, RT-qPCR assays, and standardization of amplification curves.

A.T. contributed to study conceptualization, data curation, funding acquisition, writing, and manuscript review.

C.M.V participated in study conceptualization, data curation, coordination of the research team, writing, and manuscript review.

## Ethics Statement

The hiPSCs (GM23279A) are commercially available at the NIGMS Human Genetic Cell Ethical approvals were attained by the ethics committee of Copa D’Or Hospital (CAAE number 60944916.5.0000.5249, approval number 1.791.182). Access to Brazilian genetic heritage was registered in the National System for the Management of Genetic Heritage and Associated Traditional Knowledge (SISGEN), under registration number AC1D978.

## Supplementary figures

**Supplementary Figure 1.**
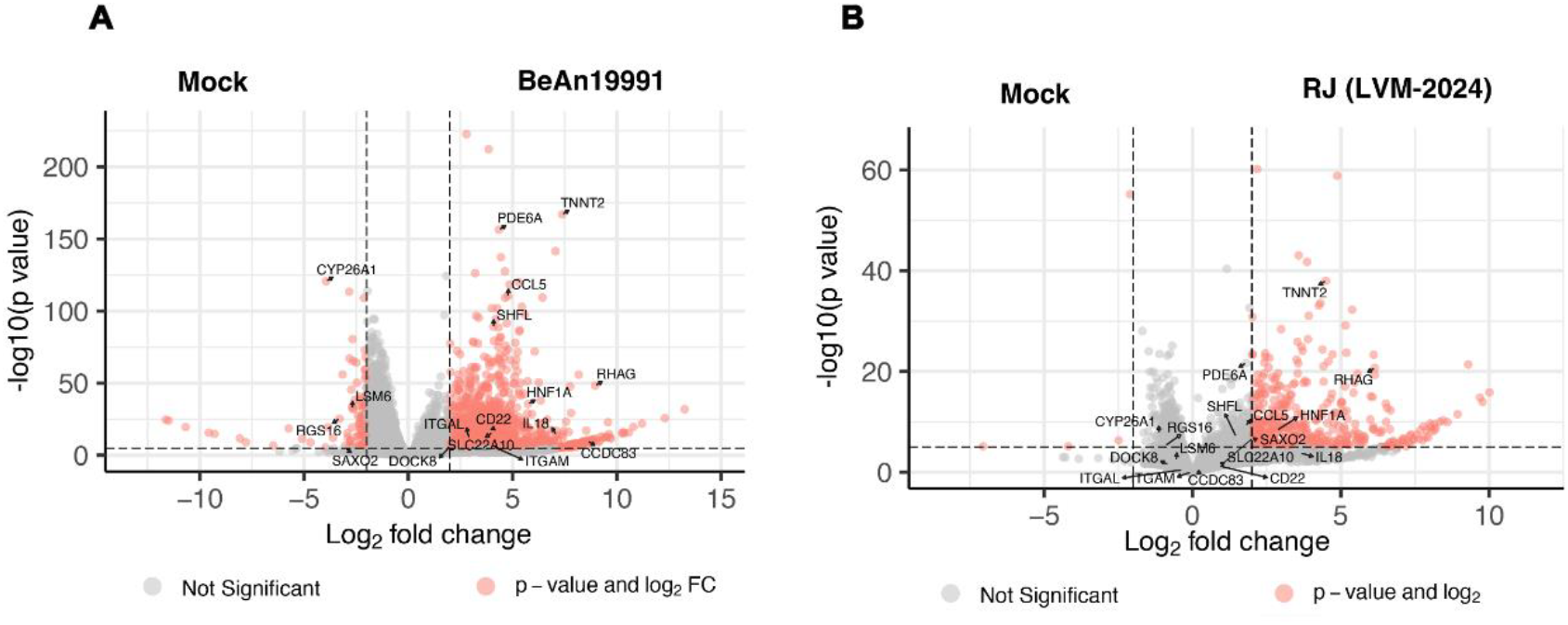
Differentially Expressed Genes between RJ (LVM-2024) and BeAn19991 against Mock. (A) Volcano plot of BeAn19991 vs Mock and (B) Volcano plot of RJ (LVM-2024) vs Mock. Differentially expressed genes (absolute Log2FoldChange higher than 2 and p value lower than 0.01) are colored in red.

**Supplementary Figure 2.**
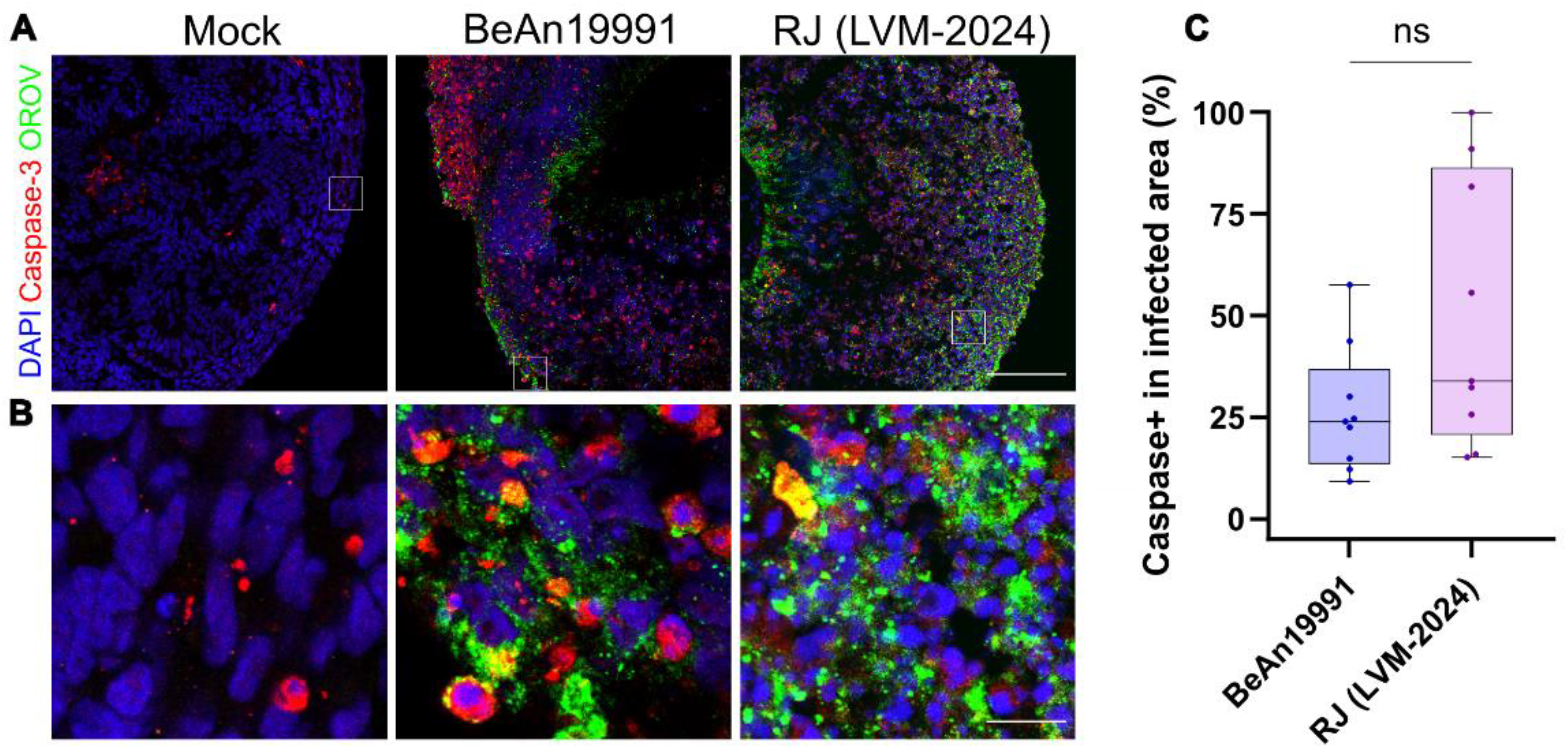
Caspase-3 activation is comparable between OROV strains. (A) Confocal images of 30-day brain organoids at 5 dpi with OROV strains BeAn19991 or RJ (LVM-2024), stained for DAPI (blue, nuclei), cleaved Caspase-3 (red, apoptotic cells), and OROV (green, viral protein). Scale bars = 100 µm. (B) High-magnification confocal images of 30-day brain organoids at 5 dpi, from the insert area in (A). Scale bars=10µm. (C) Quantification of Caspase-3^+^ in infected area in 30-day organoids at 5 dpi. Data represent mean ± SEM from two independent experiments, with at least 3 organoids in each condition.

**Supplementary Table S1. Differential expression analysis of protein-coding genes in response to OROV infection**. Results of differential gene expression analysis (DESeq2) comparing three experimental conditions using only protein-coding genes. For each comparison: (i) BeAn19991 *vs Mock*, (ii) RJ (LVM-2024) *vs Mock*, and (iii) RJ (LVM-2024) *vs* BeAn19991.

## References

1 Zhang Y, Liu X, Wu Z, et al. Oropouche virus: A neglected global arboviral threat. Virus Res 2024; 341: 199318.

2 Benitez AJ, Alvarez M, Perez L, et al. Oropouche Fever, Cuba, May 2024. Emerg Infect Dis 2024; 30: 2155–9.

3 Gräf T, Delatorre E, Ferreira C do N, et al. Expansion of Oropouche virus in nonendemic Brazilian regions: analysis of genomic characterisation and ecological drivers. The Lancet Infectious Diseases 2025; 25: 379–89.

4 Naveca FG, Almeida TAP de, Souza V, et al. Human outbreaks of a novel reassortant Oropouche virus in the Brazilian Amazon region. Nat Med 2024; 30: 3509–21.

5 Moreira FRR, Monteiro FLL, de Menezes MT, et al. Genomic evidence of Oropouche virus autochthonous circulation in a small district in the state of Rio de Janeiro, Brazil. Microbiology Spectrum 2025; 13: e02850–24.

6 Bandeira AC, Pereira FM, Leal A, et al. Fatal Oropouche Virus Infections in Nonendemic Region, Brazil, 2024. Emerg Infect Dis 2024; 30: 2370–4.

7 Schwartz DA. Novel Reassortants of Oropouche Virus (OROV) Are Causing Maternal-Fetal Infection During Pregnancy, Stillbirth, Congenital Microcephaly and Malformation Syndromes. Genes (Basel) 2025; 16: 87.

8 Garcez PP, Loiola EC, Madeiro da Costa R, et al. Zika virus impairs growth in human neurospheres and brain organoids. Science 2016; 352: 816–8.

9 Musso D, Gubler DJ. Zika Virus. Clin Microbiol Rev 2016; 29: 487–524.

10 Rodrigues AH, Santos RI, Arisi GM, et al. Oropouche virus experimental infection in the golden hamster (Mesocrisetus auratus). Virus Res 2011; 155: 35–41.

11 Santos RI, Bueno-Júnior LS, Ruggiero RN, et al. Spread of Oropouche Virus into the Central Nervous System in Mouse. Viruses 2014; 6: 3827–36.

12 Martins FE das N, Chiang JO, Nunes BTD, et al. Newborns with microcephaly in Brazil and potential vertical transmission of Oropouche virus: a case series. The Lancet Infectious Diseases 2025; 25: 155–65.

13 Lancaster MA, Renner M, Martin C-A, et al. Cerebral organoids model human brain development and microcephaly. Nature 2013; 501: 373–9.

14 Kim J, Koo B-K, Knoblich JA. Human organoids: model systems for human biology and medicine. Nat Rev Mol Cell Biol 2020; 21: 571–84.

15 Porciúncula LO, Goto-Silva L, Ledur PF, Rehen SK. The Age of Brain Organoids: Tailoring Cell Identity and Functionality for Normal Brain Development and Disease Modeling. Front Neurosci 2021; 15. DOI:10.3389/fnins.2021.674563.

16 Depla JA, Mulder LA, de Sá RV, et al. Human Brain Organoids as Models for Central Nervous System Viral Infection. Viruses 2022; 14: 634.

17 Johnson KN, Zeddam J-L, Ball LA. Characterization and Construction of Functional cDNA Clones of Pariacoto Virus, the First Alphanodavirus Isolated outside Australasia. Journal of Virology 2000; 74: 5123–32.

18 Naveca FG, Nascimento VA do, Souza VC de, Nunes BTD, Rodrigues DSG, Vasconcelos PF da C. Multiplexed reverse transcription real-time polymerase chain reaction for simultaneous detection of Mayaro, Oropouche, and Oropouche-like viruses. Mem Inst Oswaldo Cruz 2017; 112: 510–3.

19 Patro R, Duggal G, Love MI, Irizarry RA, Kingsford C. Salmon provides fast and bias-aware quantification of transcript expression. Nat Methods 2017; 14: 417–9.

20 Goto-Silva L, Ayad NME, Herzog IL, et al. Computational fluid dynamic analysis of physical forces playing a role in brain organoid cultures in two different multiplex platforms. BMC Developmental Biology 2019; 19: 3.

21 Obermeier B, Daneman R, Ransohoff RM. Development, maintenance and disruption of the blood-brain barrier. Nat Med 2013; 19: 1584–96.

22 Aguiar RS, Pohl F, Morais GL, et al. Molecular alterations in the extracellular matrix in the brains of newborns with congenital Zika syndrome. Sci Signal 2020; 13: eaay6736.

23 Ministério da Saúde. Painel Epidemiologico Oropouche. Ministério da Saúde. 2025. https://www.gov.br/saude/pt-br/assuntos/saude-de-a-a-z/o/oropouche/painel-epidemiologico/painel-epidemiologico-oropouche (accessed June 24, 2025).

24 Schwartz DA, Dashraath P, Baud D. Oropouche Virus (OROV) in Pregnancy: An Emerging Cause of Placental and Fetal Infection Associated with Stillbirth and Microcephaly following Vertical Transmission. Viruses 2024; 16: 1435.

25 Moreira FRR, Dutra JVR, Carvalho AHB de, et al. Oropouche virus genomic surveillance in Brazil. The Lancet Infectious Diseases 2024; 24: e664–6.

26 Scachetti GC, Forato J, Claro IM, et al. Re-emergence of Oropouche virus between 2023 and 2024 in Brazil: an observational epidemiological study. Lancet Infect Dis 2025; 25: 166–75.

27 Ribas Freitas AR, Schwartz DA, Lima Neto AS, Rodrigues R, Cavalcanti LPG, Alarcón-Elbal PM. Oropouche Virus (OROV): Expanding Threats, Shifting Patterns, and the Urgent Need for Collaborative Research in Latin America. Viruses 2025; 17: 353.

28 Megli CJ, Zack RK, McGaughey JJ, et al. Oropouche virus infects human trophoblasts and placenta explants. Nat Commun 2025; 16: 6040.

29 Almeida GM, Souza JP, Mendes ND, et al. Neural Infection by Oropouche Virus in Adult Human Brain Slices Induces an Inflammatory and Toxic Response. Front Neurosci 2021; 15. DOI:10.3389/fnins.2021.674576.

30 Qian X, Song H, Ming G-L. Brain organoids: advances, applications and challenges. Development 2019; 146: dev166074.

